# TOMOMAN: a software package for large scale cryo-electron tomography data preprocessing, community data sharing, and collaborative computing

**DOI:** 10.1101/2024.05.02.589639

**Authors:** Sagar Khavnekar, Philipp S. Erdmann, William Wan

## Abstract

Cryo-electron tomography (cryo-ET) and subtomogram averaging (STA) are becoming the preferred methodologies for investigating subcellular and macromolecular structures in native or near-native environments. While cryo-ET is amenable to a wide range of biological problems, these problems often have data processing requirements that need to be individually optimized, precluding the notion of a one-size-fits-all processing pipeline. Cryo-ET data processing is also becoming progressively more complex due to an increasing number of packages for each processing step. Though each package has its own strengths and weaknesses, independent development and different data formats makes them difficult to interface with one another. TOMOMAN (TOMOgram MANager) is an extensible package for streamlining the interoperability of packages, enabling users to develop project-specific processing workflows. TOMOMAN does this by maintaining an internal metadata format and wrapping external packages to manage and perform preprocessing, from raw tilt-series data to reconstructed tomograms. TOMOMAN can also export this metadata between various STA packages. TOMOMAN also includes tools for archiving projects to data repositories; allowing subsequent users to download TOMOMAN projects and directly resume processing where it was previously left off. By tracking essential metadata, TOMOMAN streamlines data sharing, which improves reproducibility of published results, reduces computational costs by minimizing reprocessing, and enables distributed cryo-ET projects between multiple groups and institutions. TOMOMAN provides a way for users to test different software packages to develop processing workflows that meet the specific needs of their biological questions and to distribute their results with the broader scientific community.

## 1. Introduction

Cryo-electron tomography (cryo-ET) is emerging as the method of choice for determining the structures of biological macromolecules *in situ*, that is, within their native context inside cells or intact extracellular particles (Young & Villa, 2023; McCafferty *et al*., 2024). Unlike other cryo-electron microscopy (cryo-EM) approaches such as single particle analysis (SPA), cryo-ET provides direct 3D reconstructions of specific fields of view, allowing one to resolve individual molecules in thick, complex specimens that would otherwise be overlapping in 2D projections. In cryo-ET, the sample is physically tilted in the microscope and projections are acquired at discreet tilt angles, resulting in a tilt-series that contains multiple distinct views of the same region of interest; these tilt-series can then be used to reconstruct 3D tomographic volumes. Subsequent high-resolution structure determination can be performed using subtomogram averaging (STA), an approach analogous to SPA methods in cryo-EM, where repeating units in the tomographic data can be extracted, aligned, averaged, and classified (Wan & Briggs, 2016; Pyle & Zanetti, 2021; Castaño-Díez & Zanetti, 2019).

Going from tilt-series data acquisition to reconstructing a tomogram involves a large number of image processing tasks (Fig. 1). We refer to this part of cryo-ET processing workflow as “preprocessing”, which occurs prior to downstream analysis such as STA. Preprocessing starts with motion correction followed by the assembly of the tilt-series in an ascending order of acquisition tilt angles. Manual curation of this tilt-series is then performed to remove bad images such as those with poor tracking, drift, or specimen charging. Curated tilt-series are then exposure filtered to reduce the high-resolution noise in later images. These exposure-filtered tilt-series must then be aligned using either fiducial based, or fiducial-less approaches. Prior to tomogram reconstruction, the contrast transfer function (CTF) needs to estimated; this can occur directly after tilt-series curation or after tilt-series alignment to make use of the alignment information. Various tomographic reconstructions can then be generated, such as raw, CTF-corrected, or denoised tomograms. The choice of which type of tomogram to reconstruct is determined by the use case; CTF-corrected tomograms provide high-resolution information for STA while denoised tomograms provide improved contrast, which can be useful for direct visual analysis.

**Figure 1.**
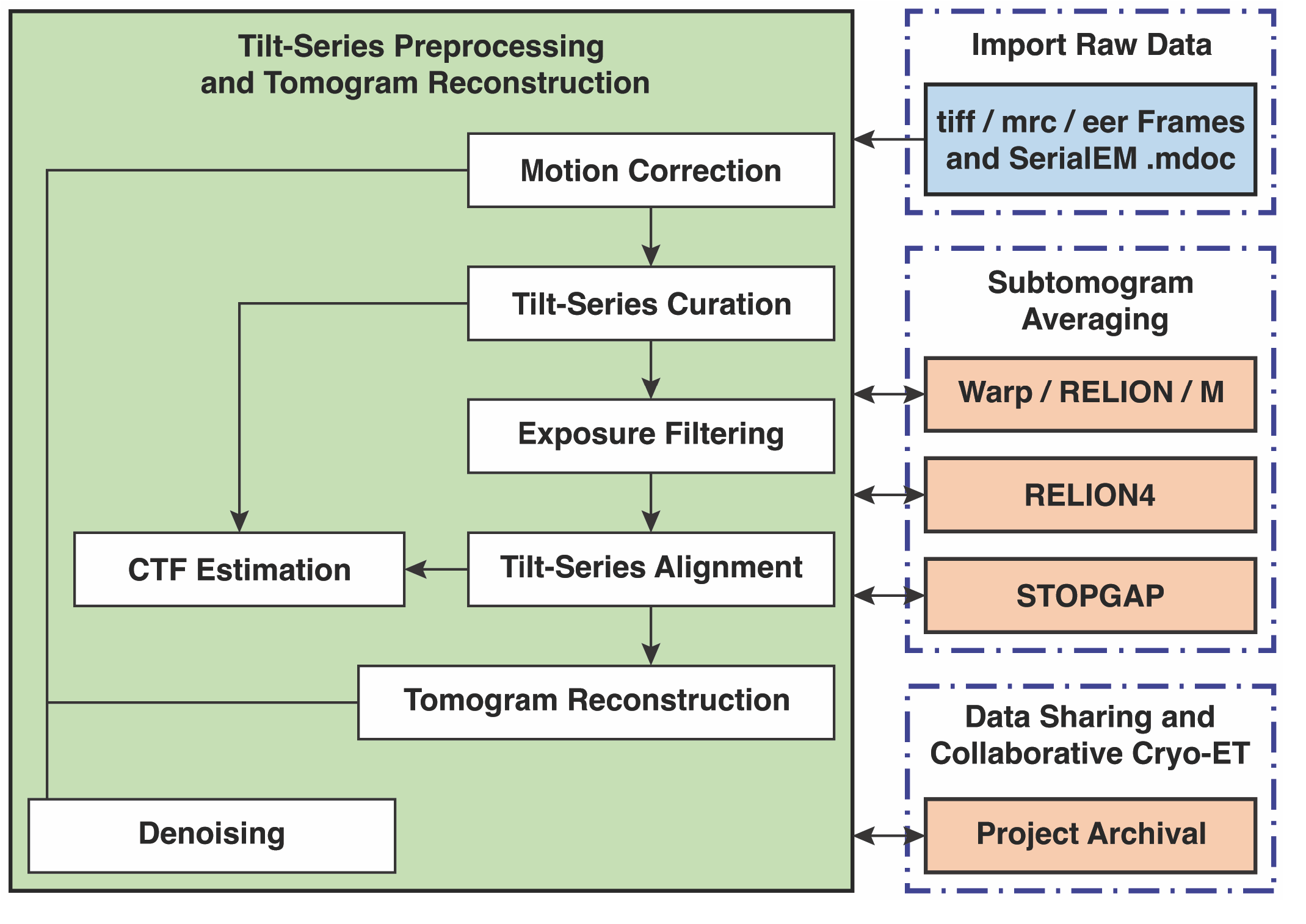
An example TOMOMAN workflow. TOMOMAN imports raw tilt-series frames and corresponding .mdoc files (top right). Preprocessing is highlighted in the light green box and individual TOMOMAN modules are depicted as white boxes; data flow is indicated by arrows. Preprocessed tilt-series and reconstructed tomograms can then be exported to subsequent STA workflows (center right), including STOPGAP, Warp/RELION/M, RELION4. Any given state of the TOMOMAN project can be archived and facilitate collaborative cryo-ET (bottom right).

Accurate preprocessing determines the success of all downstream analysis, spurring an increasing number of new approaches for each of these preprocessing tasks. These packages are often developed by different groups with different design philosophies, often with different inputs, outputs, and file formats. As such, combining various packages to tailor workflows for specific biological problems is often complex and requires significant user effort.

TOMOMAN (Tomogram Manager) is an extensible MATLAB-based package developed to reduce the complexity of combining different packages in order to streamline the testing and development of cryo-ET preprocessing and STA workflows. TOMOMAN achieves this by maintaining the essential metadata in an HDF database file (MATLAB structure format) for each tilt-series in the dataset. This metadata includes information such as filenames and paths, data collection parameters, preprocessing parameters such as estimated defocus, and tilt-series alignment information. TOMOMAN mainly acts as a wrapper for external packages, storing relevant metadata in its internal format and managing the input, output, and running of external packages. This includes managing resources for parallel processing in high performance computing (HPC) environments. As part of its design ethos, TOMOMAN is meant to be pipeline agnostic, where we make no assumption of an ideal or standard processing workflow. TOMOMAN is designed to help interface the various software packages used in each step or task of cryo-ET preprocessing so that users can test and determine the best workflow for their specific biological projects.

For subsequent STA, various pieces of preprocessing metadata are required, though the specific types and formatting of this metadata is different for each STA package. As with preprocessing, TOMOMAN facilitates the export of tilt-series data and preprocessing metadata into the correct format for each STA package, such as STOPGAP (Wan *et al*., 2024), RELION (Zivanov *et al*., 2022; Bharat & Scheres, 2016), and Warp/M (Tegunov & Cramer, 2019; Tegunov *et al*., 2021). Furthermore, TOMOMAN can also facilitate the transfer of projects between these STA packages to enable users to make use of their specific strengths.

As the cryo-ET field expands and an increasing amount of experimental data is collected, it is becoming virtually impossible for a single group to possess the requisite human effort, computational resources, and biological expertise to fully analyze all the information within the data-rich tomograms. As such, there is a need for simultaneous collaborative efforts to fully leverage this data to yield novel biological insights; these efforts can take the form of inter-institutional collaborations or larger research consortia. Furthermore, such collaborative efforts are still unlikely be able to completely analyze all the biological information, making open data archival a requirement. Current practices of depositing either raw frame data, partially processed data such as tilt-series, or final reconstructed tomograms are not sufficient as they do not include all the necessary information to reproduce published results or provide consistency to others that perform additional preprocessing. Furthermore, repeating preprocessing tasks using the same methods becomes a significant waste of resources. Maintaining consistency between independent preprocessing efforts is paramount for building richly-annotated, accessible data that provides biological insights to the broader community beyond structural biologists. Beyond streamlining cryo-ET workflows, TOMOMAN also enables reproducible data sharing and archival.

## 2. The TOMOMAN Workflow

### 2.1 An overview of TOMOMAN

TOMOMAN is an open-source package written in MATLAB that can be run in three different ways. First is as a MATLAB toolbox that can be sourced and directly used in MATLAB with a license. The second is a pre-compiled standalone package, which allows for interactive command-line use using the freely available MATLAB Compiler Runtime, similar to the Dynamo subtomogram averaging package (Castaño-Díez *et al.*, 2012; Castaño-Díez, 2017). The third way is a pre-compiled “parallel” executable, which can be used in high-performance parallel computing environments using workload managers such as SLURM (Yoo *et al.*, 2003). In addition to managing metadata, the TOMOMAN parallel executable also transparently manages the computational resource usage of external packages.

At the core of TOMOMAN is the internal metadata format and a well-defined directory structure (Fig. 2). Within each project’s root directory, a subdirectory is generated for each tilt-series in the dataset; all subsequent preprocessing tasks are performed within these subdirectories. Each tilt-series directory contains a subdirectory with the raw frames, the raw non-motion-corrected tilt stack if available, and the corresponding SerialEM-formatted tilt-series acquisition metadata (i.e. the .mdoc file) (Mastronarde, 2005). For each preprocessing step, the intermediate input and output data is stored in a subdirectory named for the software package used. The relevant metadata generated by each preprocessing step is parsed and stored in an HDF-formatted file in the project root directory; this file is referred to as the “tomolist”. Reconstructed tomograms are not considered essential data, as they are often very large and can be readily reconstructed from the annotated metadata faster than they can be transmitted. As such, tomograms are stored in their own directories separate from the tilt-series directories.

**Figure 2.**
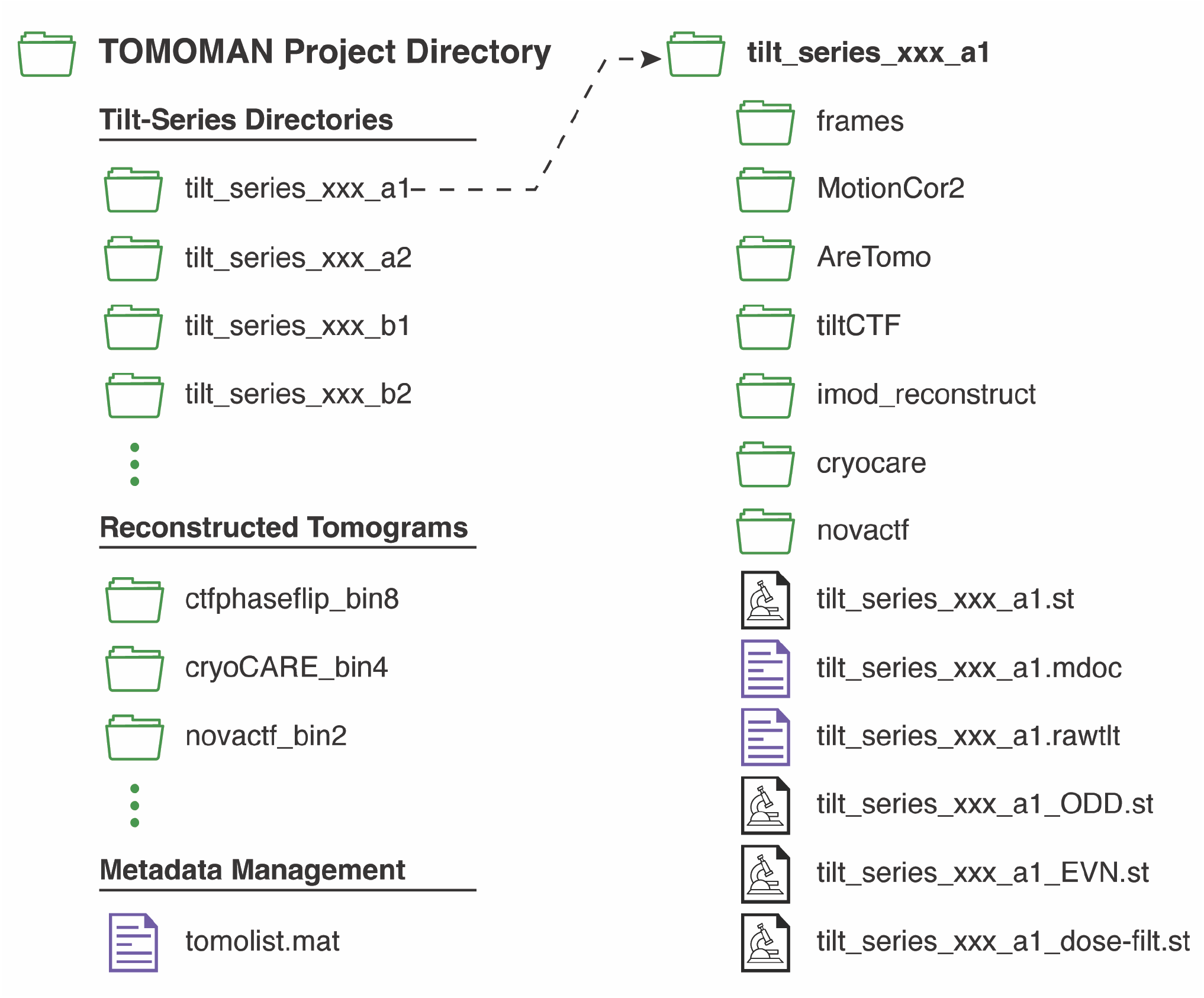
An example TOMOMAN project directory structure. The main project directory includes subfolders for each tilt-series, reconstructed tomograms, and metadata files such as the tomolist. Each tilt-series subdirectory includes further subdirectories for raw tilt-series frames and individual preprocessing tasks. Tilt-series directories also contain original microscope metadata in serial-EM formats, motion-corrected and curated tilt-series, and corresponding odd and even frame tilt-series, and dose-filtered stacks, if performed.

Below are descriptions of each preprocessing task performed in TOMOMAN. Each of these tasks is configured using package-specific TOMOMAN parameter files, which are a set of plain-text files that include the parameters for each preprocessing step as name-value pairs. Parameters generally include several TOMOMAN specific parameters, and for external packages, each input parameter for that package. TOMOMAN tasks are programmed as modules within the package, allowing for the addition of new software packages to accomplish pre-existing tasks or the development of new preprocessing tasks. Currently supported packages will be described in the documentation of each TOMOMAN release.

### 2.2 Importing microscope data

The first step in TOMOMAN preprocessing is importing raw microscope data and sorting them into subdirectories for each tilt-series. During this step, TOMOMAN scans a designated raw data directory for .mdoc files. For each one it finds, it generates the tilt-series directory. For new projects, TOMOMAN also initializes the tomolist; repeated importing will append the previous tomolist with new data, allowing for continuous TOMOMAN preprocessing during data acquisition.

### 2.3 Generating motion-corrected tilt-series

After raw microscope data has been imported, frames need to be motion corrected and subsequent aligned images assembled into a tilt-series. TOMOMAN includes modules for performing motion correction using either MotionCor2 (Zheng *et al*., 2017) or using RELION’s implementation of MotionCor2 (Zivanov *et al*., 2019). After motion correction, TOMOMAN assembles the summed frame-aligned images into tilt-series by tilt-angle, accounting for the tilt acquisition scheme used (Hagen *et al*., 2017). TOMOMAN can also output odd and even frame stacks for later use in noise2noise-based denoising (Buchholz *et al*., 2018).

### 2.4 Curating and cleaning tilt-series data

After assembling motion-corrected tilt-series, manual curation of tilt images is often necessary to remove “bad” images. Bad images include those where something blocks the field of view, such as a grid bar or ice crystal, or image quality is poor due to issues such as sample charging or drift. TOMOMAN facilitates this curation process using the *clean_stacks* task, which opens each tilt-series in IMOD’s 3dmod program (Kremer *et al*., 1996), requests input on bad images, and removes the bad images from tilt-series while annotating this in the tomolist. Tilt-series that are completely bad can also be noted in the tomolist, which then removes them from further preprocessing tasks.

### 2.5 Exposure filtering

Exposure filtering (Grant & Grigorieff, 2015) significantly improves the contrast, alignment quality, and high-resolution signals of tilt-series and reconstructed tomograms (Schur *et al*., 2016; Wan & Briggs, 2016). TOMOMAN has a module for exposure filtering on a per-tilt or per-frame basis using the empirical values determined by Grant and Grigorieff. Frame-based exposure filtering keeps track of cumulative electron exposure and uses a modified normalization filter; frame-based filtering also requires aligned frame stacks, which can be generated by MotionCor2.

### 2.6 Tilt-series alignment

The next step in the preprocessing workflow is to determine the tilt-series alignment parameters. There are two approaches to performing tilt-series alignment: fiducial-based alignment where distinct objects such as gold nanoparticles are applied to the specimen and tracked as points across the tilt-series, or fiducial-less approaches that make direct use of the image data. TOMOMAN includes modules for performing tilt-series alignment using either IMOD (Mastronarde & Held, 2017) or AreTomo (Zheng *et al*., 2022).

TOMOMAN’s *imod_preprocess* module uses the IMOD etomo package to perform initial tasks of coarse alignment and generation of necessary files for various steps within IMOD’s tilt-series alignment workflow. Users can use either etomo’s fiducial based or fiducial-less (patch tracking) approach. After initial steps are performed, IMOD’s tilt-series alignment workflow requires manual curation steps, which are performed outside of TOMOMAN in the etomo GUI.

For fully-automated fiducial-less tilt-series alignment, a module for AreTomo is also included. TOMOMAN also offers additional functionality on top of AreTomo by providing additional parameters such as user-defined binning of tilt-series prior to alignment, the use of unfiltered or exposure-filtered tilt-series as inputs, and per-tomogram thickness values. We find that these additions can be helpful when tuning parameters for specific datasets.

### 2.7 CTF estimation

As with single particle images, high-resolution STA requires the estimation and correction of the CTF. For CTF estimation, TOMOMAN includes modules for CTFFIND4 (Rohou & Grigorieff, 2015). for which CTF estimation is independent of tilt-series alignment.

Additionally, TOMOMAN includes tiltCTF, an algorithm we developed that uses the determined tilt-series alignment parameters to generate power spectra that account for the tilt-dependent defocus gradient. These power spectra are then supplied to CTFFIND4 for Thon ring fitting.

### 2.8 Tomogram reconstruction

After tilt-series alignment, TOMOMAN can use the determined alignment parameters to reconstruct tomograms using different algorithms (Fig. 3), depending on the needs of the user. Tomograms reconstructed using weighted back projection (WBP) without CTF-correction are typically output by the tilt-series alignment software (Fig 3A); these can be useful for quick visualization or low-resolution STA, but are otherwise limited in use.

**Figure 3.**
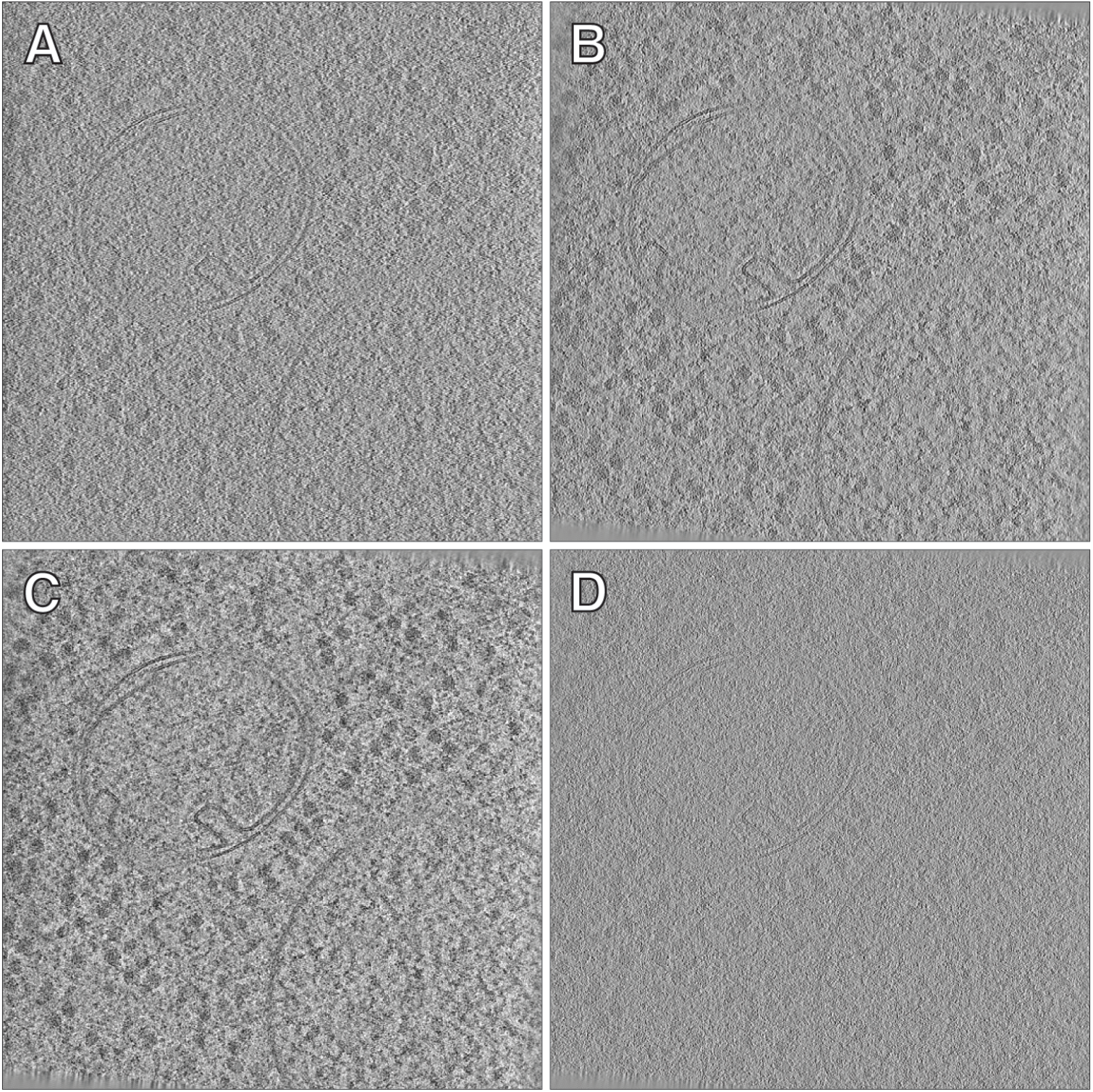
Comparison of tomograms reconstructed with different algorithms. A) Non-CTF-corrected tomogram using WBP in AreTomo. This is a tomogram reconstructed during tilt-series alignment. B) denoised with cryo-CARE (Tomograms used for training and inference are reconstructed with WBP in IMOD using odd and even frames). C) using 15 iterations of a SIRT-like filter. D) Tomogram reconstructed with 3D-CTF correction and WBP using novaCTF.

TOMOMAN can also reconstruct contrast-enhanced tomograms, which can be useful for visual analysis (Fig. 3B,C). For denoising using noise2noise methods such as cryoCARE (Fig. 3B) (Buchholz *et al*., 2018), TOMOMAN can generate tomograms using odd and even sums of motion corrected frames, which are necessary for these algorithms. Options for generating the required odd and even frame sums and corresponding tilt-series are included as parameters during the motion correction task (see section 2.3). Additionally, TOMOMAN can also reconstruct tomograms in IMOD using WBP and the SIRT-like (Simultaneous Iterative Reconstructive Technique) filter (Fig. 3C).

After tilt-series alignment and CTF estimation, CTF-corrected tomograms can be reconstructed. TOMOMAN also includes a module to generate 2D CTF-corrected tomograms using IMOD’s tilt and ctfphaseflip programs (Xiong *et al*., 2009) as well as 3D CTF-correction using novaCTF (Turoňová *et al*., 2017). For novaCTF, TOMOMAN will generate the appropriate scripts as well as temporary and output directory for running novaCTF. After reconstruction, TOMOMAN will also perform any desired tomogram binning by Fourier cropping using the Fourier3D program (Turoňová, 2020). CTF-corrected tomograms can then be used for subsequent high-resolution STA.

### 2.9 Tomogram denoising

It is becoming a common practice to use machine learning methods to enhance the contrast in tomograms through denoising. Contrast enhanced or denoised tomograms allow for easier interpretation of molecular features by the human eye. TOMOMAN supports generation of denoised tomograms using using cryoCARE (Buchholz *et al*., 2018). TOMOMAN handles generating tomograms from odd and even frames (see above) and handles the generation of necessary training files and the execution cryoCARE.

### 2.10 Automated pipelines

While all the above tasks can be run individually, it is often convenient to run as many tasks as possible in an automated workflow, particularly when preprocessing large high-throughput datasets. TOMOMAN allows users to define the various tasks of their preprocessing workflows into a “pipeline”, i.e. a list of preprocessing tasks, which can then be run on high-performance computing clusters.

### 2.11 Interoperability with STA workflows

In addition to the core pre-processing workflow described in the previous section, TOMOMAN includes additional tools to export TOMOMAN metadata to other STA workflows. These include directly exporting to STOPGAP (Wan *et al*., 2024), Warp-RELION3-M (Tegunov & Cramer, 2019; Bharat & Scheres, 2016; Tegunov *et al*., 2021), and RELION4 (Zivanov *et al*., 2022) as well as moving between each of these workflows. For the Warp/RELION/M workflow, TOMOMAN also handles motion corrected tilt images, while excluding those removed during curation, corresponding .mdoc files, tilt-series alignment files, and RELION 3 formatted particle list. For the RELION 4 tomography workflow, TOMOMAN handles curated tilt-series, tilt-series alignment and CTF estimation parameters, additional file with order of tilt-series acquisition, and per tomogram particle list in RELION 4 tomography star file format. For STOPGAP, to generate required metadata files, such as wedge lists, functions are provided to export them directly from the tomolist. Additionally, TOMOMAN can also convert particle metadata from STOPGAP particle list into RELION star file format.

## 3. Archival and data sharing

### 3.1 Minimal projects for archival

TOMOMAN includes an archival module to export TOMOMAN project to a so-called “minimal project” that can be deposited to repositories such as EMPIAR (Iudin *et al*., 2023). Minimal projects retain the TOMOMAN directory structure, tomolist, and only the files necessary to revive the project at its current preprocessing state at a later time point or at another location, while cleaning up unnecessary intermediate files generated during preprocessing. The ability to revive a project in its exact state is key to reproducibility, as small changes in preprocessing can significantly affect downstream results. For example, slight differences in fiducial centerring causes differences in tilt-series alignment and subsequent tomogram reconstruction; this affects the particle positions and orientations within tomograms as well as the resolution of STA structures. This minimizes the size of the project directory, as it is only necessary to keep the raw frames, motion-corrected tilt stacks, and the metadata and parameter files, such as estimated CTF parameters and tilt-series alignment parameters. This 2D imaging data is typically only a total of 2 – 3 gigabytes in size, while metadata and parameter files are on the order of kilobytes. This data is sufficient to reconstruct tomograms for subsequent STA workflows. This reduction in size makes it easier to share data, as large files such as tomograms, which can be hundreds of gigabytes, can often be reconstructed faster than they can be transmitted.

### 3.2 Community data sharing and distributed collaborative cryo-ET

TOMOMAN minimal projects allow users to effectively restore projects to their exact previous preprocessing states. In addition to being key to reproducibility and archival, this also enables distribution of preprocessed projects between collaborators for more complex downstream processing such as STA. This enables large-scale cellular cryo-ET projects, where different labs can focus on their specific molecules or subcellular structures of interest.

## 4. Conclusion

One major challenge in cryo-ET and STA is how to leverage the unique capabilities of the wide variety of packages available for each step of the image processing workflow. This is typically due to the difficulty in managing the metadata between these packages, which typically have different file and parameter formats. TOMOMAN addresses these issues by establishing its own internal metadata format and providing extensible modules to interface with other software packages, allowing users to develop their own package-agnostic workflows that are best suited to their biological problems.

TOMOMAN is an open-source package written in MATLAB that is supplied as a standalone package that can be executed using the freely available MATLAB runtime. TOMOMAN is designed with high performance computing in mind; it generates all necessary scripts to launch external packages with defined computational resources, allowing users to seamlessly run custom workflows in parallel. Beyond the initial cryo-ET preprocessing tasks, TOMOMAN’s metadata tracking enables transferring projects between different STA workflows including Warp/RELION3/M, RELION4, and STOPGAP.

Other cryo-ET packages such as IMOD (Kremer *et al*., 1996), Warp (Tegunov & Cramer, 2019), EMAN2 (Tang *et al*., 2007; Chen *et al*., 2019), ScipionTomo (Jiménez de la Morena *et al*., 2022),TomoBEAR (Balyschew *et al*., 2023), and RELION5 also offer start to end workflows for tilt-series preprocessing. However, these packages generally aim to offer a simplified, streamlined workflow that may not be suitable for each biological problem. IMOD, Warp and EMAN2 primarily use their own internal functions for preprocessing tasks and tomographic reconstruction, while RELION5 now includes a cryo-ET workflow that provides wrapper scripts for external packages to perform CTF estimation, tilt-series alignment, denoising and an internal tomogram reconstruction algorithm. ScipionTomo on the other hand offers a workflow manager where individual packages can be added as plugins. TomoBEAR, is similar in functionality to TOMOMAN in that it wraps external packages and runs scripted workflows, but it is aimed at minimizing intermediate steps to deliver a streamlined, linear, STA workflow. TOMOMAN also streamlines the interoperability of various packages, but aims to facilitate the testing and use of different packages for each preprocessing step. This flexibility allows users to optimize workflows to their specific biological problems.

One unique feature of TOMOMAN is its archival functions, which streamline data deposition to community databases such as EMPIAR while also providing the necessary metadata for users to download datasets and restart projects. This enables the sharing of information-rich cryo-ET datasets without the need for downstream users to reprocess data, thereby reducing overall computational costs and ensuring reproducibility between labs. This aspect of reproducibility is particularly important, as subtle changes in preprocessing can significantly affect downstream results. We believe these archival functions will help enable large-scale consortium cryo-ET projects, while also opening the data to the wider biological community. TOMOMAN has already been used to manage a number of projects including some that have been deposited to EMPIAR as TOMOMAN minimal projects such as EMPIAR-11830 (Khavnekar *et al*., 2023), EMPIAR-11756 (Khavnekar *et al*., 2023), EMPIAR-11658 (Wan *et al*., 2024; Rangan *et al*., 2023), EMPIAR-11398 (Khavnekar *et al*., 2022), EMPIAR-11325, EMPIAR-11324, and EMPIAR-11322 (Khavnekar, *et al*., 2023). In particular, the EMPIAR-11830 project was also processed as a multi-institutional, multi-user collaborative TOMOMAN project. We envision that these TOMOMAN minimal project depositions together with well-annotated metadata will be important for the development of large data approaches, such as novel AI-based image processing tools.

Altogether, TOMOMAN offers a solution for developing comprehensive cryo-ET workflows that are accessible for both experienced and novice users. The software package is available at https://github.com/wan-lab-vanderbilt/TOMOMAN.

## Acknowledgements

This work was supported by U.S. National Institutes of Health grant DP2GM146321 (to WW). WW is a Pew Scholar in the Biomedical Sciences, supported by the Pew Charitable Trusts. This work was conducted in part using the resources of the Advanced Computing Center for Research and Education at Vanderbilt University, Nashville, TN. Some of this work was performed at the Max Planck Institute of Biochemistry with support from Wolfgang Baumeister, Juergen Plitzko, and John Briggs for resources and infrastructure. We would also like to thank Max Planck Computing and Data Facility (MPCDF) for the computational infrastructure. SK would like to thank the Max Planck Society and the International Max Planck Research School for graduate school funding.

## References

Balyschew, N., Yushkevich, A., Mikirtumov, V., Sanchez, R. M., Sprink, T. & Kudryashev, M. (2023). Nat Commun 14, 6543.

Bharat, T. A. M. & Scheres, S. H. W. (2016). Nat Protoc 11, 2054–2065.

Buchholz, T.-O., Jordan, M., Pigino, G. & Jug, F. (2018). *arXiv* 10.48550/arXiv.1810.05420.

Castaño-Díez, D. (2017). Acta Cryst D 73, 10.1107/S2059798317003369.

Castaño-Díez, D., Kudryashev, M., Arheit, M. & Stahlberg, H. (2012). Journal of Structural Biology 178, 139–151.

Castaño-Díez, D. & Zanetti, G. (2019). Current Opinion in Structural Biology 58, 68–75.

Chen, M., Bell, J. M., Shi, X., Sun, S. Y., Wang, Z. & Ludtke, S. J. (2019). Nat Methods 16, 1161–1168.

Grant, T. & Grigorieff, N. (2015). Elife 4, e06980.

Hagen, W. J. H., Wan, W. & Briggs, J. A. G. (2017). Journal of Structural Biology 197, 191–198.

Iudin, A., Korir, P. K., Somasundharam, S., Weyand, S., Cattavitello, C., Fonseca, N., Salih, O., Kleywegt, G. J. & Patwardhan, A. (2023). Nucleic Acids Research 51, D1503–D1511.

Jiménez de la Morena, J., Conesa, P., Fonseca, Y. C., de Isidro-Gómez, F. P., Herreros, D., Fernández-Giménez, E., Strelak, D., Moebel, E., Buchholz, T. O., Jug, F., Martinez-Sanchez, A., Harastani, M., Jonic, S., Conesa, J. J., Cuervo, A., Losana, P., Sánchez, I., Iceta, M., del Cano, L., Gragera, M., Melero, R., Sharov, G., Castaño-Díez, D., Koster, A., Piccirillo, J. G., Vilas, J. L., Otón, J., Marabini, R., Sorzano, C. O. S. & Carazo, J. M. (2022). Journal of Structural Biology 214, 107872.

Khavnekar, S., Kelley, R., Waltz, F., Wietrzynski, W., Zhang, X., Obr, M., Tagiltsev, G., Beck, F., Wan, W., Briggs, J., Engel, B., Plitzko, J. & Kotecha, A. (2023). Microscopy and Microanalysis 29, 961–963.

Khavnekar, S., Vrbovská, V., Zaoralová, M., Kelley, R., Beck, F., Klumpe, S., Kotecha, A., Plitzko, J. & Erdmann, P. S. (2022). *bioRxiv* 2022.06.16.496417.

Khavnekar, S., Wan, W., Majumder, P., Wietrzynski, W., Erdmann, P. S. & Plitzko, J. M. (2023). Journal of Structural Biology 215, 107911.

Kremer, J. R., Mastronarde, D. N. & McIntosh, J. R. (1996). Journal of Structural Biology 116, 71–76.

Mastronarde, D. N. (2005). Journal of Structural Biology 152, 36–51.

Mastronarde, D. N. & Held, S. R. (2017). J Struct Biol 197, 102–113.

McCafferty, C. L., Klumpe, S., Amaro, R. E., Kukulski, W., Collinson, L. & Engel, B. D. (2024). Cell 187, 563–584.

Pyle, E. & Zanetti, G. (2021). Biochemical Journal 478, 1827–1845.

Rangan, R., Khavnekar, S., Lerer, A., Johnston, J., Kelley, R., Obr, M., Kotecha, A. & Zhong, E. D. (2023). *bioRxiv* 2023.08.18.553799.

Rohou, A. & Grigorieff, N. (2015). Journal of Structural Biology 192, 216–221.

Schur, F. K. M., Obr, M., Hagen, W. J. H., Wan, W., Jakobi, A. J., Kirkpatrick, J. M., Sachse, C., Kräusslich, H.-G. & Briggs, J. A. G. (2016). Science 353, 506–508.

Tang, G., Peng, L., Baldwin, P. R., Mann, D. S., Jiang, W., Rees, I. & Ludtke, S. J. (2007). Journal of Structural Biology 157, 38–46.

Tegunov, D. & Cramer, P. (2019). Nat Methods 16, 1146–1152.

Tegunov, D., Xue, L., Dienemann, C., Cramer, P. & Mahamid, J. (2021). Nat Methods 18, 186–193.

Turoňová, B. (2020). https://github.com/turonova/Fourier3D.

Turoňová, B., Schur, F. K. M., Wan, W. & Briggs, J. A. G. (2017). Journal of Structural Biology 199, 187–195.

Wan, W. & Briggs, J. a. G. (2016). Methods Enzymol 579, 329–367.

Wan, W., Khavnekar, S. & Wagner, J. (2024). Acta Cryst D 80, 10.1107/S205979832400295X.

Xiong, Q., Morphew, M. K., Schwartz, C. L., Hoenger, A. H. & Mastronarde, D. N. (2009). J Struct Biol 168, 378–387.

Yoo, A. B., Jette, M. A. & Grondona, M. (2003). Vol. Job Scheduling Strategies for Parallel Processing, edited by D. Feitelson, L. Rudolph & U. Schwiegelshohn. pp. 44–60. Berlin, Heidelberg: Springer.

Young, L. N. & Villa, E. (2023). Annual Review of Biophysics 52, 573–595.

Zheng, S. Q., Palovcak, E., Armache, J.-P., Verba, K. A., Cheng, Y. & Agard, D. A. (2017). Nat Methods 14, 331–332.

Zheng, S., Wolff, G., Greenan, G., Chen, Z., Faas, F. G. A., Bárcena, M., Koster, A. J., Cheng, Y. & Agard, D. A. (2022). Journal of Structural Biology: X 6, 100068.

Zivanov, J., Nakane, T. & Scheres, S. H. W. (2019). IUCrJ 6, 5–17.

Zivanov, J., Otón, J., Ke, Z., von Kügelgen, A., Pyle, E., Qu, K., Morado, D., Castaño-Díez, D., Zanetti, G., Bharat, T. A., Briggs, J. A. & Scheres, S. H. (2022). eLife 11, e83724.

